# PPIAT: Target Mass Spectrometry-based Protein-Protein Interaction Analytics Tool

**DOI:** 10.1101/2023.04.04.535369

**Authors:** Jongham Park, Ahrum Son, Hyunsoo Kim

## Abstract

**Motivation:** The formation of protein networking is critical for understanding the biological functions of proteins and the underlying mechanisms of disease. To gain insights into target proteins that interact with a particular disease, we need to profiling all the proteins involved in the disease beforehand. However, the profiling results may not necessarily match with an experimental target protein. Therefore, it is necessary to identify physical protein-protein interactions (PPIs) using various methods. The cross-linking mass spectrometry (XL-MS) method is a representative approach to identify physical interactions between proteins, but there are many challenges in calculating theoretical mass values for application to target mass spectrometry.

**Results:** The research team developed PPIAT, which integrates information on reviewed human proteins, protein-protein interactions, cross-linkers, enzymes, and modifications. All functions are available for free on the web application.

**Availability and implementation:** The PPIAT is available at www.ppiat.cnu.ac.kr

## 1. Introduction

The formation of protein networks and protein complex structures through protein-protein interactions (PPIs)^[1-4]^ is crucial for understanding the biological functions of proteins. Many human disease result from abnormal PPIs involving endogenous proteins, proteins from pathogens or both.^[4-6]^ For example aberrant PPIs have been implicated in a various of human diseases, including cancers.^[6]^ To understand a target protein’s interaction with a specific disease, we need to profile all proteins involved in the disease beforehand. However, the profiling results may not match with an experimental target protein.^[7-9]^ Therefore, it is essential to identify physical protein-protein interactions using several methods. In addition to understanding the networking of interactions between proteins, it is also necessary to comprehend the interaction at the proteome-wide level.^[10-12]^ Search for theoretical protein-protein interaction is available in several web-based databases such as STRING^[12]^, MINT^[13]^,BioGRID^[14]^ InAct^[15]^, and the development of these resources make a significant contribution to protein interactome profiling analysis.

There are methodologies to find out these interaction sites, such as covalent labeling^[16]^ footprinting^[17]^, hydrogen-deuterium exchange mass spectrometry (HDX-MS)^[18]^, cross-linking mass spectrometry (XL-MS, sometimes abbreviated CX-MS or CL-MS)^[19-27]^, ion-mobility MS^[28]^, and native MS^[29]^.^[10, 29]^ Among these methods, XL-MS has been developed over several decades and has become a powerful method for mapping protein-protein interactions.^[24]^ However, its effectiveness has long been impeded by three primary obstacles: (1) complex tandem mass spectrometry (MS/MS or MS2) fragmentation of cross-linked peptides; (2) low abundance of cross-linked peptides in complex peptide mixtures; (3) heterogeneity of cross-linked products;^[10]^ (4) It is very difficult to calculate all possible cases and their mass values by considering the site, charge, and modification where interactions between proteins actually occur; (5) The first hurdle makes accurate identification of cross-linked peptides and unambiguous assignment of cross-linked sites difficult, the second and third hurdles hinder effective MS detection of cross-linked peptides.^[4]^ The fourth hurdle becomes one of the major factors hindering the efficiency of research. Target mass spectrometry (MRM or SRM) methods generally used to identify PPIs, it performs based on MS/MS values obtained through profiling. But MS/MS fragmentation of cross-linked peptides is typically convoluted and unpredictable. Therefore, to identify the site of the predicted theoretical actual interaction occurs and calculate their mass values, we need to compare experimental MS/MS spectra against a computed library of theoretical spectra. This process of calculating mass values must not only consider all cases of theoretical interaction but also the properties of cross-linker, charge of peptide/fragmented peptide ion, and modification in this process of calculating mass values. Finally, there are some tools to help analyze XL-MS like SRMcollider^[30]^, Prego^[31]^ and X-Link Transition Calculator^[32]^ in skyline. The SRMcollider is a software which predict interference probability between target transitions.^[30]^ The Prego is a software which predicts high responding peptides for SRM experiments.^[31]^ The X-Link Transition Calculator is a software which calculate cross linked mass value.^[32]^ In general, targeted methods perform based on MS/MS values obtained through profiling. However, these tools are unable to search theoretical protein interactors for the target protein, and it must be known which sites are actual interact and modified already. In this respect, development of analytics tools for XL-MS is still needed.

We suggest PPIAT: a target mass spectrometry-based protein-protein interaction analytics tool. It is a web-based analytics tool that can search interaction information about human proteins and calculate mass values given the properties of cross-linkers, enzymes, charge, and modifications. We expect PPIAT to improve experimental efficiency and become a significant contributor to cater to theoretical information for XL-MS analysis.

## 2. Materials and methods

### 2.1 Overview of PPIAT

PPIAT was built using XAMPP, CI3 (Codeignator3), and CSS (Cascading Style Sheet). XAMPP includes the Apache web server, PHP (Hypertext Preprocessor), and MySQL, so the server runs on Apache, and the database was built based on MySQL. PPIAT has three main functions. First, it allows users to search for human proteins by utilizing information on reviewed human proteins from UniProt.^[25][33]^ Second, it enables users to search for information about commercialized cross-linkers, including their name, binding site, cleavability^[26]^, spacer arm length, and mass values. The mass values are differentiated between non-cleavage mass values and cleavage mass values. Third, PPIAT can predict protein-protein interactions for the user’s target protein and calculate their mass values, considering input values such as target protein, cross-linker, enzyme, peptide length range, peptide charge, ion charge, modifications on the search page. Finally, the results calculated from PPIAT can be summarized and extracted based on the probability score of protein interaction in the derived results and other analysis tools such as Prego^[31]^. Through these functions, PPIAT can be used as a tool for targeted mass spectrometry-based protein interactor analysis in XL-MS. (Figure. 1)

**Fig. 1.**
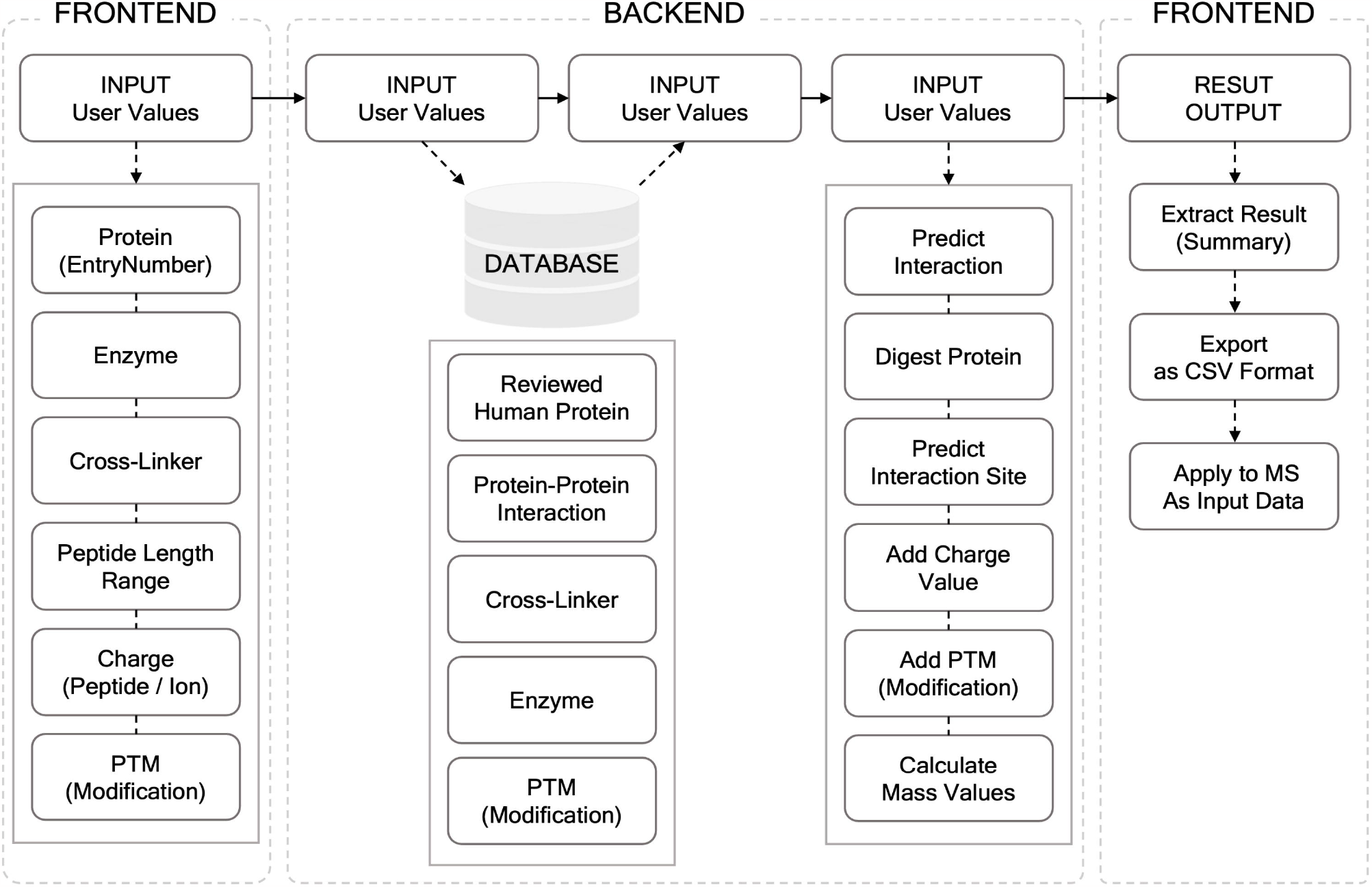
Workflow of PPIAT the figure presents the flow of the PPIAT. Information for searching proteinprotein interaction (PPIs) is input at front-end. Data queried from database by input information at front-end and PPIs and each mass value are calculated considering input condition. All calculated data presents at front-end and the result can export as CSV format. The exported data can use as input data for MS/MS analysis.

### 2.2 Building database

PPIAT integrates data from reviewed human proteins, protein-protein interactions, cross-linkers, enzymes, and modifications. It includes 20,386 reviewed human protein data (43,277 including isoforms), around 20 billion interaction data, and 83 cross-linker data. These data were collected from UniProt^[33]^, STRING^[12]^, and publicly published white papers. The unreviewed human protein data is approximately ten times larger than the reviewed human protein data. However, as the data on unreviewed human protein is unverified, it has not been considered.

### 2.3 User Guide about PPIAT

PPIAT is easy to use with several input values, including (1) the entry number of the target protein that the user wants to search, (2) the enzyme that digests the input target protein, (3) the cross-linker that is utilized in XL-MS^[19-27]^ to capture protein-protein interactions, (4) the peptide length range that the user wants to check at the MS/MS step, (5) the peptide charge and fragmented peptide ion charge, and (6) the modification that is predicted to occur in the protein-protein interactome. (see Figure 2).

**Fig. 2.**
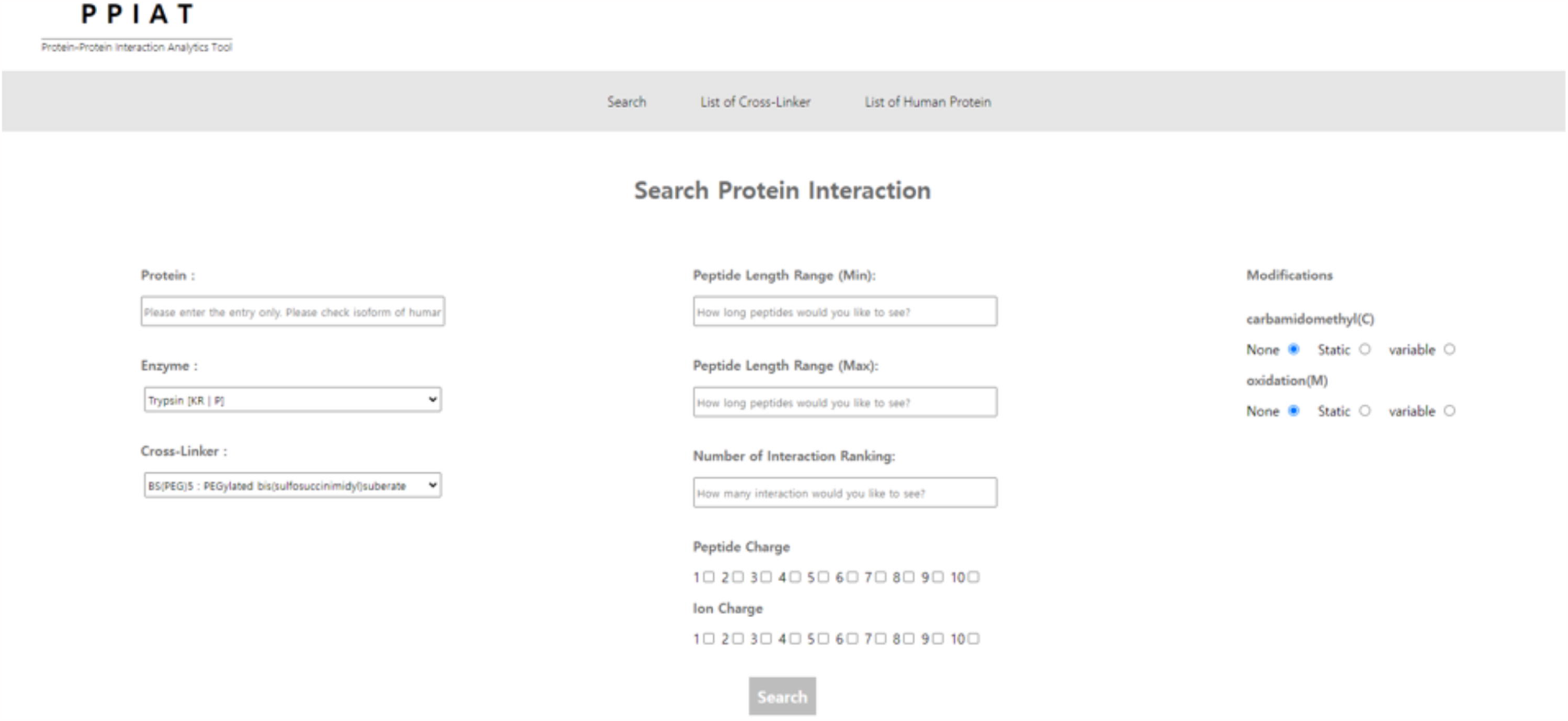
Search page of PPIAT This is the page for searching PPIs and their mass value. For search, there needs some information for search such as entrynumber of protein, an enzyme for protein digestion, cross-linker for cross-linking, range of peptide length, ranking of protein-protein interaction, charges of peptide and ion, and protein modification.

The information of reviewed human protein at UniProt^[33]^ includes several columns, such as ID, name, organism, entry number, entry name, and sequence ID. The entry number and entry name information can be used to search for protein-protein interactions at resources like STRING^[12]^.

Proteins are differentiated between canonical and isoform, where the isoform is a member of a set of highly similar proteins that originate from a single gene or gene family and are the result of genetic differences. They have different mass values since they have the same or similar function but not an equal sequence.^[34]^ For this reason, PPIAT divides the same entry name and protein name between canonical and isoform using a dash (“-”) in the entry number. Even though the values of the string column at UniProt^[33]^ and values of protein A, and protein B at STRING^[12]^ are linked to each other (e.g., format of 9606.ENSP00000379873), they are not totally matched, and the canonical and isoform use the same value. In other words, since the values of protein A and protein B at STRING^[12]^ are not differentiated between canonical and isoform, PPIAT cannot identify the user’s target protein as a canonical and isoform. For this reason, to distinguish between canonical and isoform, PPIAT uses the entry umber for searching the target protein.

Users can select trypsin or chymotrypsin as an enzyme that digests the target protein, considering the properties of the digestion site and exception. Additionally, users can select a cross-linker from the list of 83 commercialized cross-linkers, to identify the interaction between a target protein and other proteins given the properties of the cross-linker binding site and cleavability. (see Data S1) Lastly, by taking into account the peptide length range, the number of interaction rankings, the charge of the peptide and fragmented peptide ion, and modification, PPIAT calculates the mass values of the target protein and its protein interactome. The peptide length range is the maximum and minimum peptide length that a user wants to identify at the interaction level, and the number of interaction ranking is the number of interactable proteins that the user wants to identify. The input values of the peptide charge and fragmented peptide ion charge considered for MS/MS analysis allow for overlapping selection. There are two types of modification, which are divided into static and variable type to calculate mass values, and they also allow for overlapping selection.

Through the above process, PPIAT can identify the interactions with a target protein and calculate the actual interaction that occurs by cross-linker in the peptide and ion levels while considering the charge of the peptide and fragmented peptide ion, properties of cross-linker and enzyme, modification.

## Result

The XL-MS workflow is divided into several steps: (1) A target complex is cross-linked in solution and digested with trypsin into peptides; (2) The peptides are analyzed by liquid chromatography coupled mass spectrometry (LC–MS/MS) to obtain precursor masses and fragment masses for cross-linked peptides; (3) The fragmentation spectra of all peptides are subjected to database searching to identify cross-linked peptides;^[21]^ (4) On result page, the output of the PPIAT can be summarized and extracted based on the probability score of protein interaction in the derived results and other analysis tools such as Prego^[31]^, and can be used as input data for target mass spectrometry analysis. The combined score is divided into four groups: highest confidence (>0.9), high confidence (>0.7), medium confidence (>0.4), and low confidence (>0.15). The analysis program Prego provides a predicts list of high-responding peptides for SRM experiments.^[31]^

The output of the PPIAT software, which searches for protein-protein interactions (PPIs) and calculates their mass values, can be used as input for MS analysis. The software generates separate tables for identified PPIs and predicts interaction information, indicating which proteins are interacting with the target protein based on combined score. The other table provides printed information on the theoretical actual interaction, including combined score, the protein A’ peptide, cross linker, protein B’ peptide, precursor charge, precursor m/z, protein A’ ion, protein A’ ion type, protein B’ ion, protein B’ ion type, product charge, and product m/z.

As a result of searching for target protein through PPIAT, a list of theoretical PPIs for the target protein, the number of all theoretical cases in which actual interactions occur in the interactome, and their mass values are calculated. The theoretical PPIs list predicts which proteins interact with the target protein with scores based on STRING database.^[12]^ In each column of the tables. In each column of the tables, “Protein A” refers to the target protein inputted, and “Protein B” refers to the proteins that interact with the target protein in peptide and ion levels. The precursor m/z and product m/z columns are divided at the peptide and ion levels into the mass value of protein A linked with one side of XL, XL Mass after cleavage, and protein B linked with the other side of XL. If the user forecasts that a modification will occur statically, fluctuating mass values are added. On the other side, if the forecast is that the modification will occur variably, the result is printed as two lines, one with add one without the change in the mass values due to the modification. (see Figure 3).

**Fig. 3.**
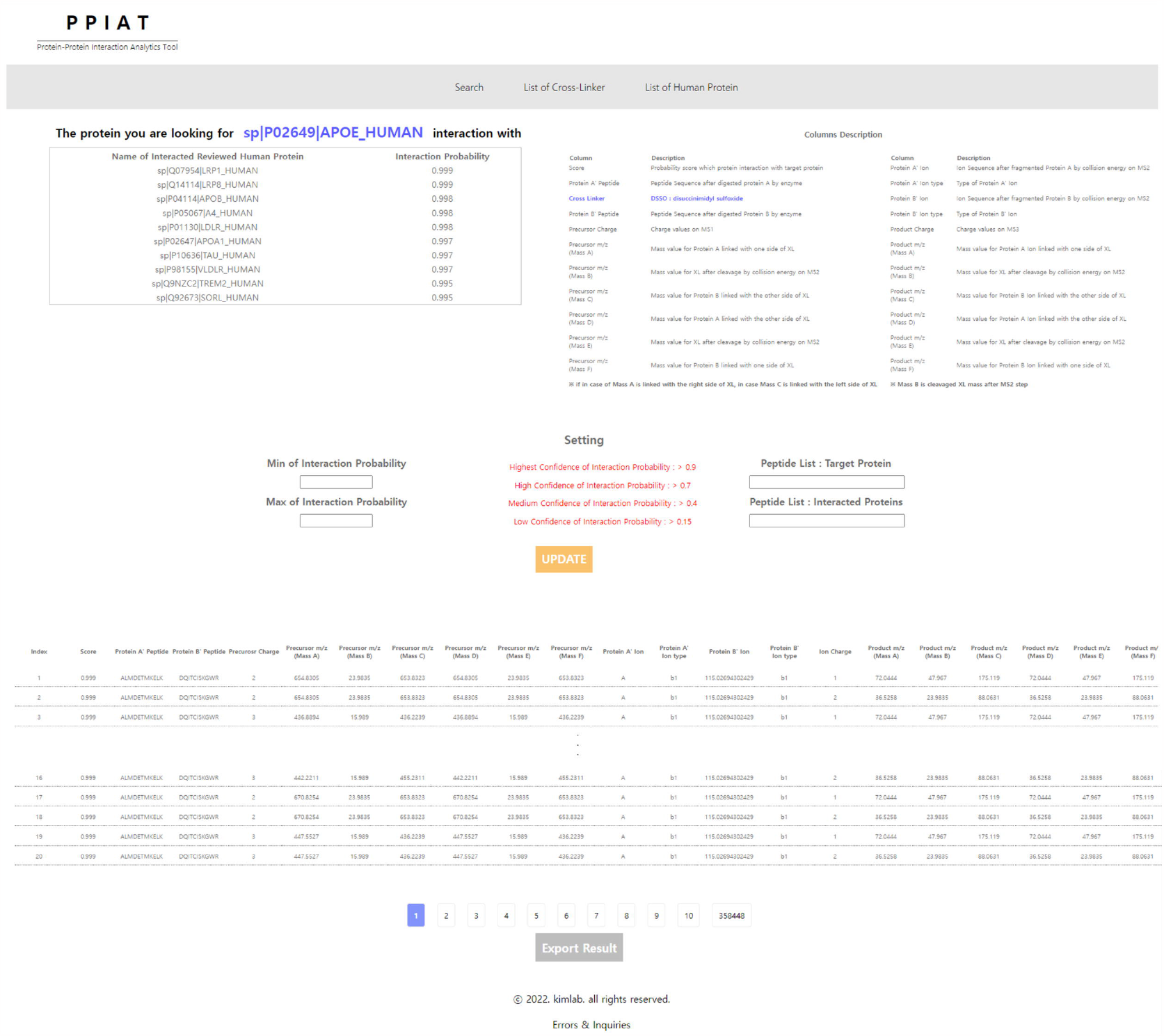
Result page at PPIAT. Output of PPIAT is consist of section about identify protein-protein interaction and calculated mass value considering condition for XL-MS.

In XL-MS method, mass spectrometry-based analysis is performed on cross-linked complexes after digestion by enzyme. PPIAT does not consider the total mass of the cross-linked complex, but only calculates the mass values of the digested cross-linking protein complexes at the peptide and fragmented peptide ion levels. For each of the proteins interacting with the target protein, it is necessary to take into account all other products of the cross-linking reaction of Protein A and Protein B interaction at the peptide and fragmented peptide ion levels: (1) cross-linked between Protein A and Protein B (cross-linking or interpeptide); (2) only cross-linked with Protein A, not cross-linked to Protein B (loop-link or intrapeptide); (3) only cross-linked with Protein B, not cross-linked to Protein A (loop-link or intrapeptide); (4) non-cross-linked between Protein A and Protein B. (mono-link or deadend).^[21,22]^ Additionally, when experimenting with XL-MS, we may encounter situations where interactions occur on one side and the other side of the cross-linker may interact with Protein A and Protein B respectively.

Cross-linkers are divided into cleavable cross-linkers and non-cleavable cross-linkers. ^[35, 36]^ Therefore, it is necessary to consider the properties of the cross-linker, the cleavage site, and how cleavage occur (e.g., symmetric or asymmetric) when using a cleavable cross-linker.^[35, 36]^ If the crosslinker to be used has a cleavable property and is cleaved symmetrically, the same mass values should be added to Protein A and Protein B, respectively. However, if the cross-linker to be used has a cleavable property and is cleaved asymmetrically, different mass values must be added. All calculated mass values can be exported to CSV format and used as input data at MS/MS analytics.

We conducted a search for the APOE E4 protein, which is a major biomarker for Alzheimer’s disease, to identify potential PPIs using the PPIAT tool. The conditions used for the search were as follows: APOE E4(UniProt ID: P02649), trypsin digestion, cross-linker DSSO, top 10 interactions, peptide length of 9, peptide charge of 2 and 3, ion charge of 1 and 2, and modifications of carbamidomethylation(static) and oxidation(variable). As a result of our search, PPIAT identified ten proteins: LRP8(UniProt ID: Q14114, score:0.999), LRP1(UniProt ID: Q07954, score:0.999), A4(APP, UniProt ID: P05067, score:0.998), APOB(UniProt ID: P04114, score:0.998), LDLR(UniProt ID: P01130, score:0.998), APOA1(UniProt ID: P02647, score:0.997), VLDLR(UniProt ID: P98155, score:0.997), TAU(MAPT, UniProt ID: P10636, score:0.997), SORL(UniProt ID: Q92673, score:0.995), and TREM2(UniProt ID: Q9NZC2, score: 0.995) (see Figure 4), along with their theoretical information in XL (see Data S2 and S3). We confirmed that the output PPIs information matched the search results from STRING. Moreover, by comparing the mass values through Skyline software, we confirmed that the mass shift was consistent with that of the cross-linker DSSO (see Data S2 and S3).

## Discussion

XL-MS is a widely used method for identifying the sites of protein interactions and elucidating the three-dimensional structure of the proteins. In recent years, significant technological advancements in XL-MS studies have propelled the proteomics field forward. However, there are still several limitations that need to be overcome. To address these challenges, various strategies have been developed, such as the use of different cross-linkers. However, there is currently no solution or analysis tool available for accurately calculating the mass values of the theoretical interactions required for MS analysis.

To address this challenge, our research team has developed PPIAT, a tool that can search for proteinprotein interactions with target proteins, identify the theoretical interactions that occur, and calculate their mass values at the peptide and fragmented peptide ion levels and accurately calculate their mass using PPIAT as input data.

Overall, the development of PPIAT represents a major step forward in XL-MS studies and has the potential to significantly enhance our understanding of protein-protein interactions. Although strategies such as the development of various cross-linkers have been used in XL-MS, there is still a lack of solutions or analysis tools for accurately calculating the mass values of the theoretical interactions required for MS analysis. To address this issue, several software tools have been developed, including SRMcollider^[30]^, Prego^[31]^, X-Link Transition Calculator^[32]^ in Skyline. By combining the three software tools above, it is possible to predict the outcome of a cross-link between proteins and utilize the results to analyze XL-MS. However, unlike PPIAT, these tools are not integrated with the platform used to search for theoretical PPIs, and therefore, the protein modification, binding site, and mass value of the cross-linker must be calculated and entered directly for XL-MS. While PPIAT is a valuable tool for XL-MS analysis, it does have some limitations that need to be addressed. Firstly, it does not cover all the products of the cross-linker, nor does it provide information about the enzyme or modification. Additionally, the output generated by PPIAT is only theoretical, and experimental verification is necessary to conform the results. To address these limitations, our team has built a database that includes information on 83 different cross-linkers, two types of enzymes (trypsin and chymotrypsin)^[21]^, and the most commonly occurring modifications (oxidation and carbamidomethylation). While these updates are a step forward, we will continue to update and improve the database on a regular basis to ensure that it remains a useful resource for the scientific community.

In the case of cross-linkers, various types with unique properties such as chemical cross-linkers and Photo/MS-cleavable cross-linkers^[35-37]^ have been studied and commercialized. Moreover, for experiments, researchers synthesize cross-linkers with novel characteristics, such as BMSO, DBB, DHSO, and SDASO, and use them.^[38-42]^ Although these synthetic cross-linkers are not widely used and therefore not included in the PPIAT database, an editing function will be added to the software to enable users to input information on cross-linkers when searching for target proteins. The PPIAT database is continuously updated with reviewed information on human proteins and protein-protein interactions. While the current version of the database has been built using PPI information from STRING^[12]^ only, we plan to expand it by integrating with other databases such as MINT^[13]^, BioGRID^[14]^, and InAct^[15]^ for PPI searching. The output generated by PPIAT can be downloaded in CSV format and used as input data for mass spectrometry. However, given the variety of MS equipment available, it is essential to format the output appropriately for each type of MS.

## Conclusions

In conclusion, this paper presents PPIAT, a web-based open platform designed for the analytics of XL-MS data. By facilitating the search for protein-protein interactions involving target proteins and calculating the mass values of all cases of theoretical interaction between proteins at both the peptide and fragmented peptide ion levels, PPIAT addresses a major hurdle in XL-MS data analysis. We anticipate that this tool will be widely adopted by researchers in the fields of proteomics and bioinformatics, and we welcome contributions to further its development. By continuing to update and refine the database, we hope to address these limitations and further enhance the accuracy and efficiency of XL-MS analysis.

## Supporting information

Table1_PPIs_Identification

DataS1_list_of_XL

DataS2_resut_of_skyline

DataS3_result_of_PPIAT

## AVAILABILITY

A source code for PPIAT are available at https://github.com/kimlab-cnu.

## ACKNOWLEDGEMENTS

Author contributions: A.R.S., and H.S.K. conceived the original idea of the project. J.H.P. developed the PPIAT analytics tool. J.H.P., A.R.S. and H.S.K. wrote the manuscript.

---

**Table 1.** APOE E4 PPIs search from PPIAT. Top 10 list of proteins interaction with APOE E4 protein. The results are matched with results on STRING.

| Rank | Protein Name | Interaction Probability |
| --- | --- | --- |
| 1 | sp\|Q07954\|LRP1_HUMAN | 0.999 |
| 2 | sp\|Q14114\|LRP8_HUMAN | 0.999 |
| 3 | sp\|P04114\|APOB_HUMAN | 0.998 |
| 4 | sp\|P05067\|A4_HUMAN | 0.998 |
| 5 | sp\|P01130\|LDLR_HUMAN | 0.998 |
| 6 | sp\|P02647\|APOA1_HUMAN | 0.997 |
| 7 | sp\|P10636\|TAU_HUMAN | 0.997 |
| 8 | sp\|P98155\|VLDLR_HUMAN | 0.997 |
| 9 | sp\|Q9NZC2\|TREM2_HUMAN | 0.995 |
| 10 | sp\|Q92673\|SORL_HUMAN | 0.995 |

