## Supplementary material for "PPIAT: Target Mass Spectrometry-based Protein-Protein Interaction Analytics Tool": DataS1_list_of_XL: Data_S1_List_of_Cross_Linker.docx

**Data S1.List of Cross linkers in PPIAT : Dashed Lines Indicate Cleavage Sites**

| Name | Cleavability | Formula | Monoisotopic | Formula  (Cleavaged) | Monoisotopic  (Cleavaged) | Structure |
| --- | --- | --- | --- | --- | --- | --- |
| BS(NHS)PEG5 | No | C_22_H_32_N_2_O_13_ | 532.19043909463 | C_22_H_32_N_2_O_13_ | 532.19043909463 | 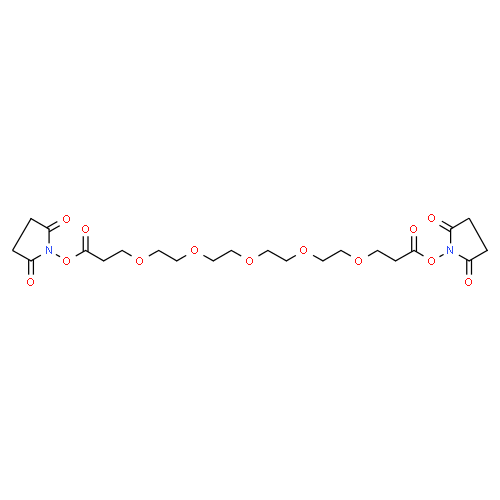 |
| BS(PEG)9 | No | C_30_H_48_N_2_O_17_ | 708.29529808859 | C_30_H_48_N_2_O_17_ | 708.29529808859 | 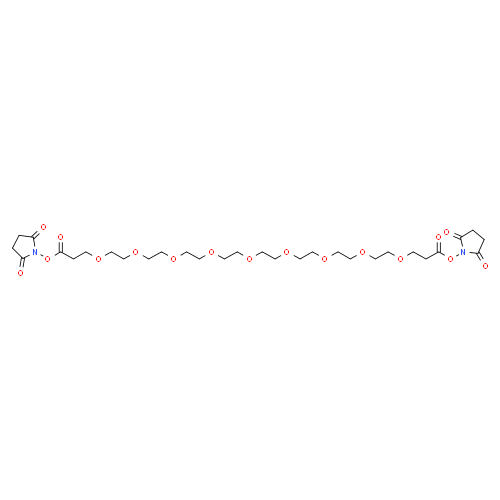 |
| BS2G-d0 | No | C_13_H_12_N_2_Na_2_O_14_S_2_ | 529.9525339824 | C_13_H_12_N_2_Na_2_O_14_S_2_ | 529.9525339824 | 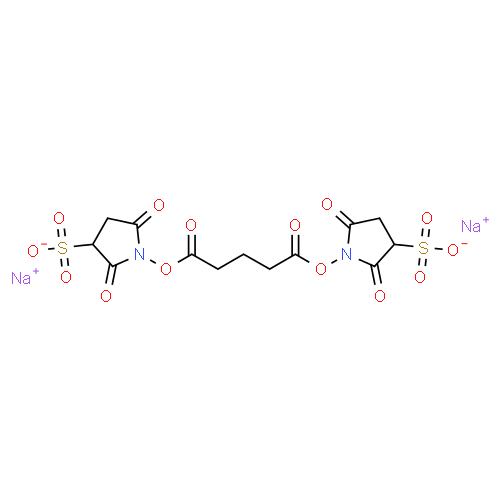 |
| BS2G-d4 | No | C_13_H_8_D_4_N_2_Na_2_O_14_S_2_ | 533.98383411132 | C_13_H_8_D_4_N_2_Na_2_O_14_S_2_ | 533.98383411132 | 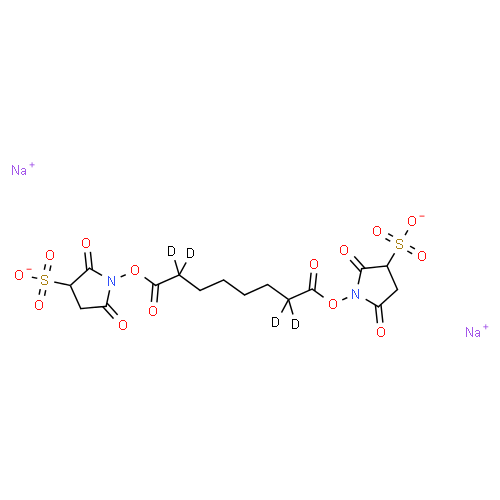 |
| BS3 | No | C_16_H_18_N_2_O_14_S_2_Na_2_ | 571.99948417578 | C_16_H_18_N_2_O_14_S_2_Na_2_ | 571.99948417578 | 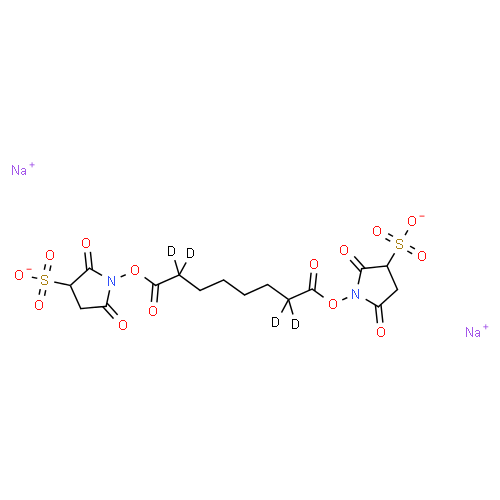 |
| BS3-d4 | No | C_16_H_14_D_4_N_2_Na_2_O_14_S_2_ | 576.0307843047001 | C_16_H_14_D_4_N_2_Na_2_O_14_S_2_ | 576.0307843047001 | 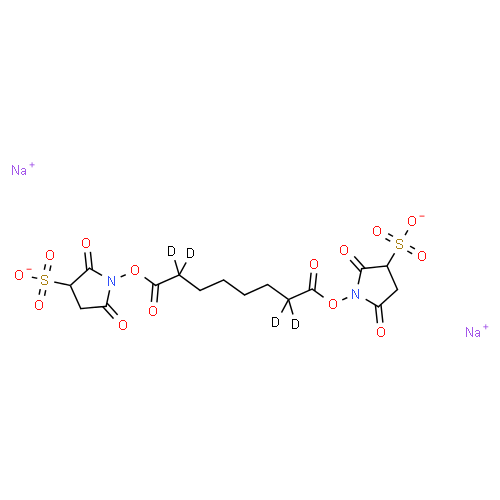 |
| DFDNB | No | C_6_H_2_F_2_N_2_O_4_ | 203.99826287706003 | C_6_H_2_F_2_N_2_O_4_ | 203.99826287706003 | 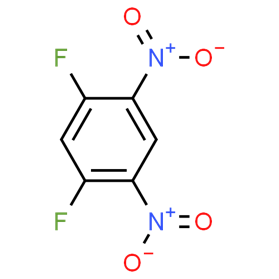 |
| DMP | No | C_9_H_20_Cl_2_N_2_O_2_ | 258.0901832566 | C_9_H_20_Cl_2_N_2_O_2_ | 258.0901832566 | 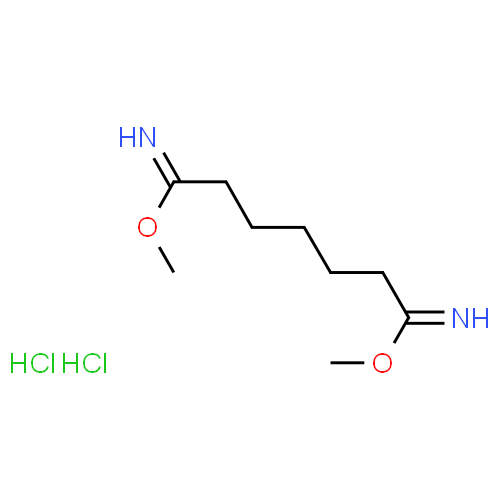 |
| DMS | No | C_10_H_20_N_2_O_2_ | 200.1524778926 | C_10_H_20_N_2_O_2_ | 200.1524778926 | 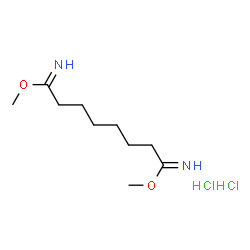 |
| DSG | No | C_13_H_14_N_2_O_8_ | 326.07501541664 | C_13_H_14_N_2_O_8_ | 326.07501541664 | 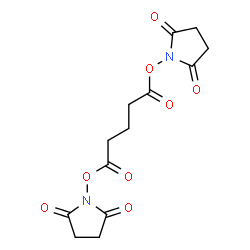 |
| DSS | No | C_16_H_20_N_2_O_8_ | 368.12196561001997 | C_16_H_20_N_2_O_8_ | 368.12196561001997 | 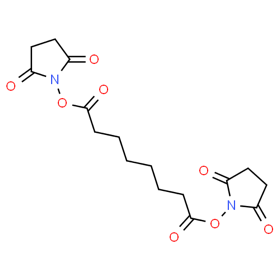 |
| TSAT | No | C_18_H_18_N_4_O_12_ | 482.09212203270005 | C_18_H_18_N_4_O_12_ | 482.09212203270005 | 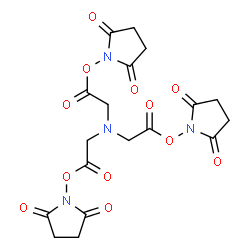 |
| DSP | Yes | C_14_H_16_N_2_O_8_S_2_ | 404.03480782990005 | C_7_H_8_NO_4_S  C_7_H_8_NO_4_S | 202.01740391495002  202.01740391495002 | 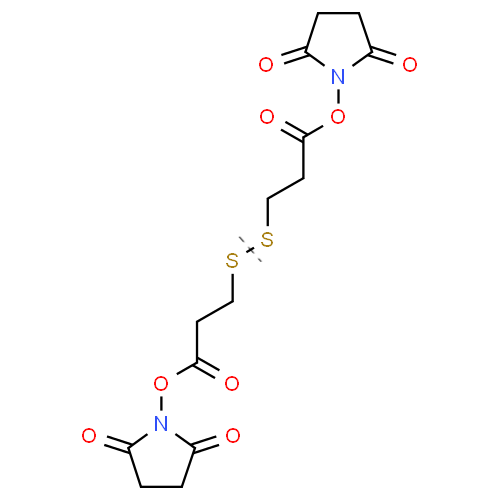 |
| DTBP | Yes | C_8_H_18_Cl_2_N_2_O_2_S_2_ | 308.01867554093997 | C_4_H_9_ClNOS  C_4_H_9_ClNOS | 154.00933777046998  154.00933777046998 | 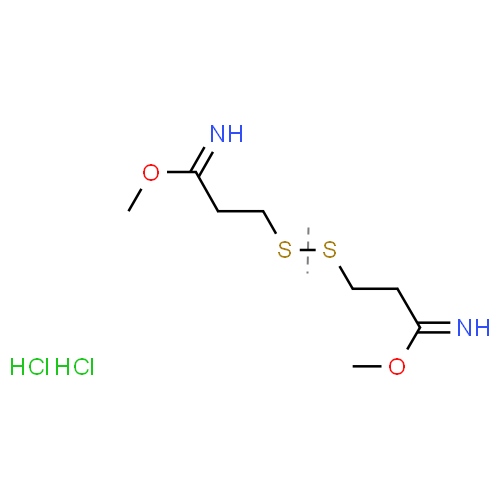 |
| DTSSP | Yes | C_14_H_14_N_2_Na_2_O_14_S_4_ | 607.9123263956601 | C_7_H_7_NNaO_7_S_2_  C_7_H_7_NNaO_7_S_2_ | 303.95616319783005  303.95616319783005 | 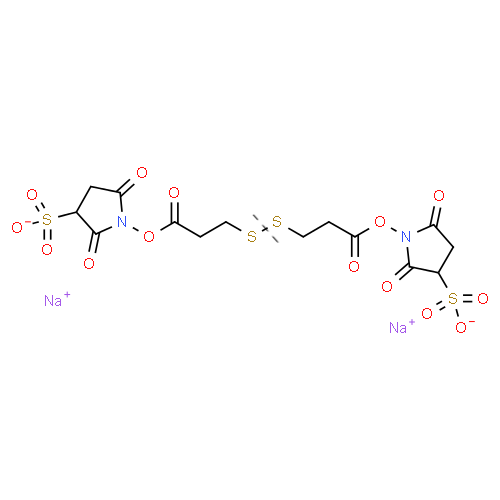 |
| EGS | Yes | C_18_H_20_N_2_O_12_ | 456.1016240883 | C_8_H_8_NO_5_  C_2_H_4_O_2_  C_8_H_8_NO_5_ | 198.04024736012002  60.02112936806  198.04024736012002 | 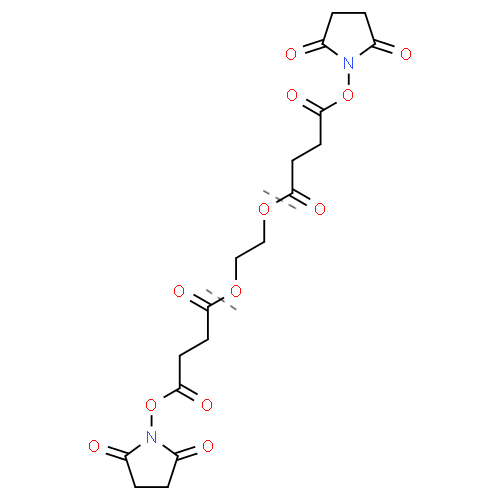 |
| Sulfo-EGS | Yes | C_18_H_18_N_2_Na_2_O_18_S_2_ | 659.9791426540601 | C_8_H_7_NNaO_8_S  C_2_H_4_O_2_  C_8_H_7_NaO_8_S | 299.979006643  60.02112936806  299.979006643 | 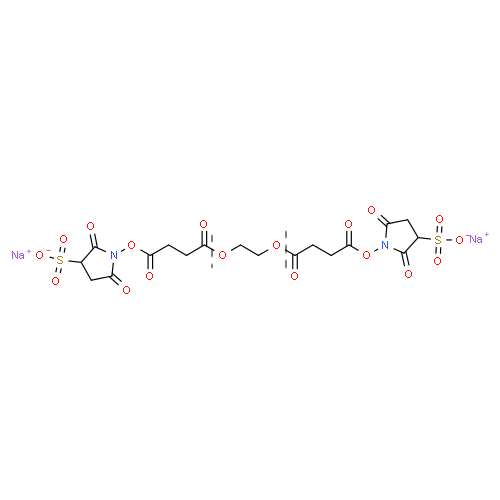 |
| DSBU | Yes | C_17_H_22_N_4_O_9_ | 426.13867830290997 | C_8_H_11_N_2_O_4_  CO  C_8_H_11_N_2_O_4_ | 199.07188184167  27.99491461957  199.07188184167 | 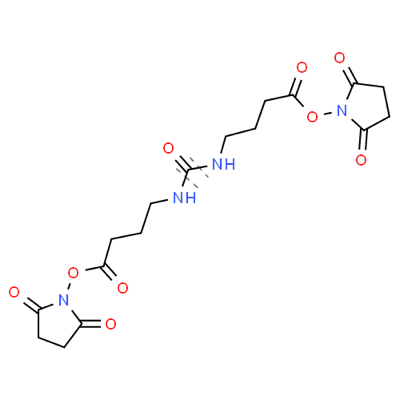 |
| DSSO | Yes | C_14_H_16_N_2_O_9_S | 388.05765127507004 | C_7_H_8_NO_4_  SO  C_7_H_8_NO_4_ | 170.04533274055  47.96698579397  170.04533274055 | 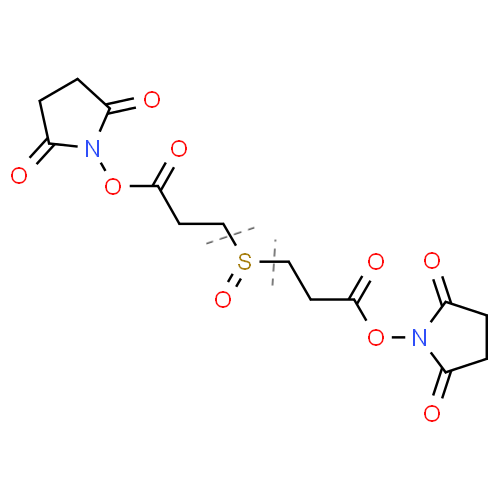 |
| DST | Yes | C_12_H_12_N_2_O_10_ | 344.04919459132 | C_6_H_6_NO_5_  C_6_H_6_NO_5_ | 172.02459729566  172.02459729566 | 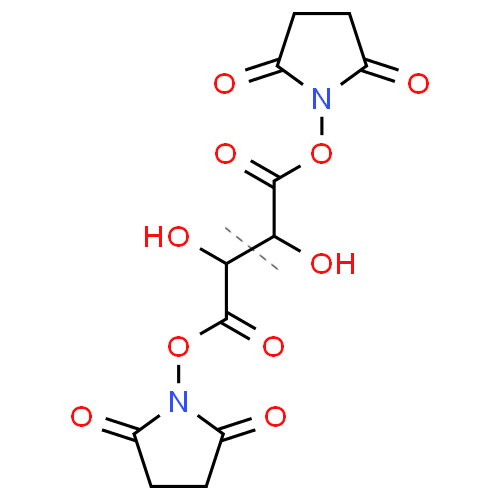 |
| NHS-Azide | No | C_6_H_6_N_4_O_4_ | 198.03890468938 | C_6_H_6_N_4_O_4_ | 198.03890468938 | 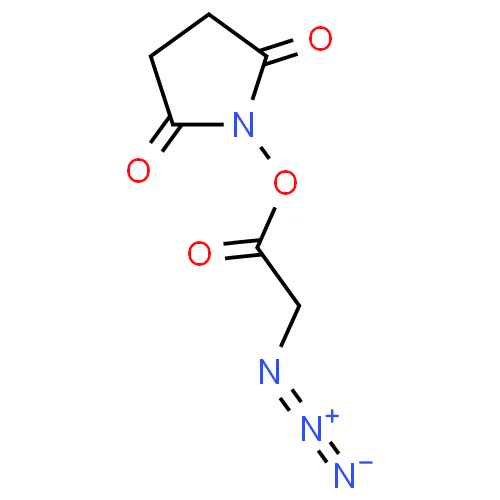 |
| NHS-PEG4-Azide | No | C_15_H_24_N_4_O_8_ | 388.15941374780004 | C_15_H_24_N_4_O_8_ | 388.15941374780004 | 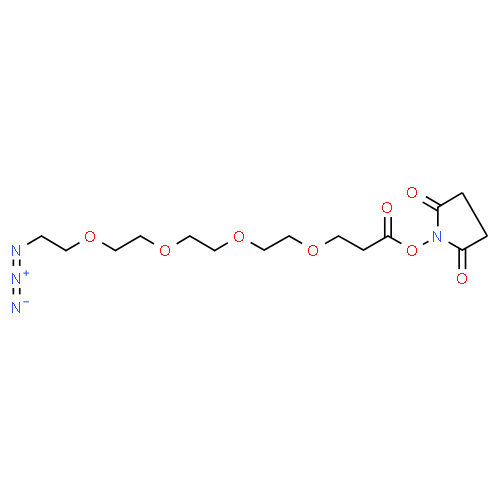 |
| NHS-Phosphine | No | C_25_H_20_NO_6_P | 461.10282436487 | C_25_H_20_NO_6_P | 461.10282436487 | 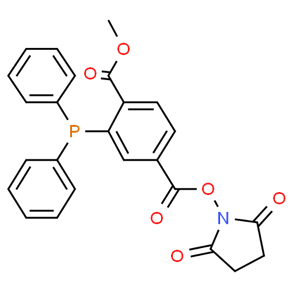 |
| EDC | No | C_8_H_18_ClN_3_ | 191.11892527543 | C_8_H_18_ClN_3_ | 191.11892527543 | 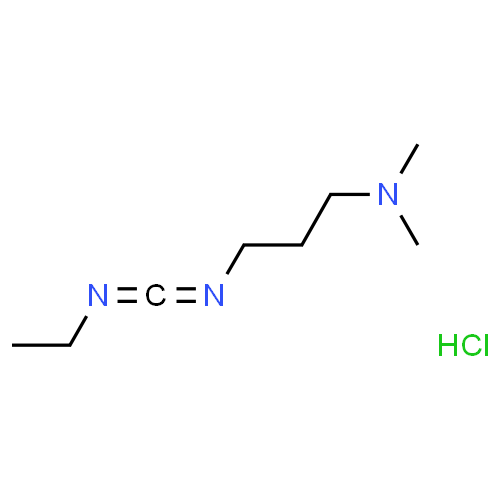 |
| NHS | No | C_4_H_5_NO_3_ | 115.02694302429 | C_4_H_5_NO_3_ | 115.02694302429 | 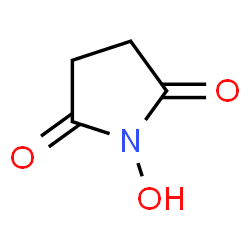 |
| Sulfo-NHS | No | C_4_H_4_NNaO_6_S | 216.96570230717003 | C_4_H_4_NNaO_6_S | 216.96570230717003 | 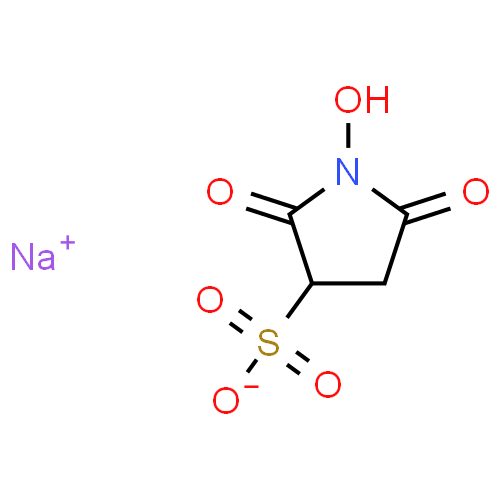 |
| LC-SDA | No | C_15_H_22_N_4_O_5_ | 338.15901982463 | C_15_H_22_N_4_O_5_ | 338.15901982463 | 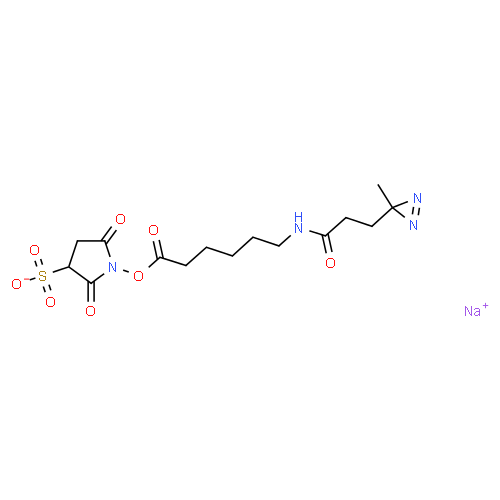 |
| SDA  (NHS-Diazirine) | No | C_9_H_11_N_3_O_4_ | 225.0749558461 | C_9_H_11_N_3_O_4_ | 225.0749558461 | 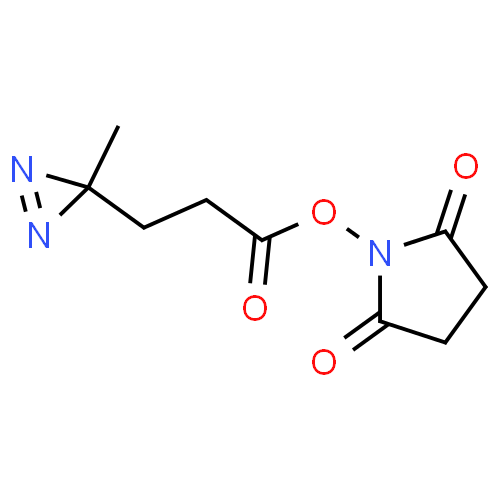 |
| SDAD  (NHS-SS-Diazirine) | Yes | C_14_H_20_N_4_O_5_S_2_ | 388.08751210897003 | C_7_H_12_N_3_OS  C_7_H_8_NO_4_S | 186.07010819402  202.01740391495002 | 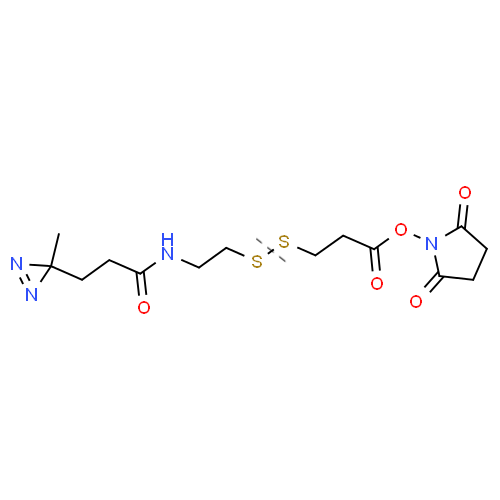 |
| SPB | No | C_19_H_15_NO_8_ | 385.07976644444 | C_19_H_15_NO_8_ | 385.07976644444 | 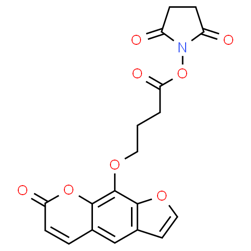 |
| Sulfo-LC-SDA | No | C_15_H_21_N_4_NaO_8_S | 440.09777910751 | C_15_H_21_N_4_NaO_8_S | 440.09777910751 | 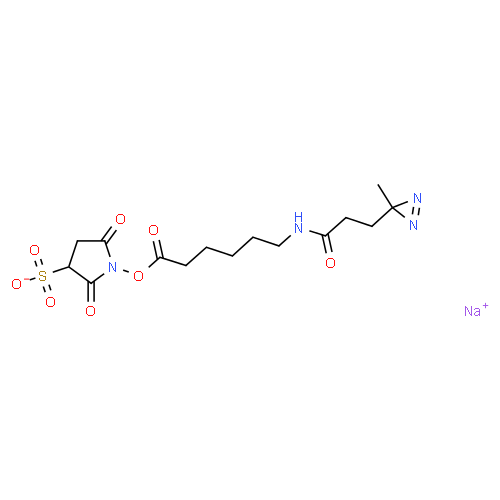 |
| Sulfo-SANPAH | No | C_16_H_17_O_9_N_6_NaS | 492.06754160702 | C_16_H_17_O_9_N_6_NaS | 492.06754160702 | 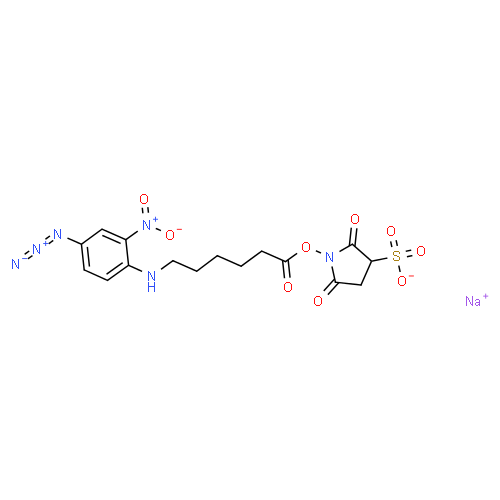 |
| Sulfo-SBED | Yes | C_32_H_42_N_9_NaO_11_S_4_ | 879.1784321884002 | C_25_H_35_N_8_O_4_S_2_  C_7_H_7_NNaO_7_S_2_ | 575.22226899057  303.95616319783005 | 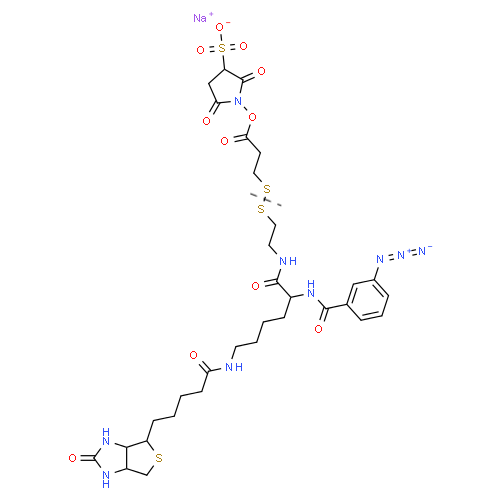 |
| Sulfo-SDA | No | C_9_H_10_N_3_Na_7_S | 352.98792848399006 | C_9_H_10_N_3_Na_7_S | 352.98792848399006 | 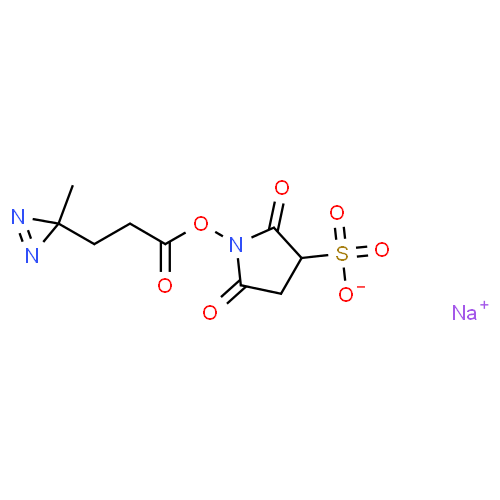 |
| Sulfo-SDAD | Yes | C_14_H_19_N_4_NaO_8_S_3_ | 490.02627139184995 | C_7_H_7_NNaO_7_S_2_  C_7_H_12_N_3_OS | 303.95616319783005  186.07010819402 | 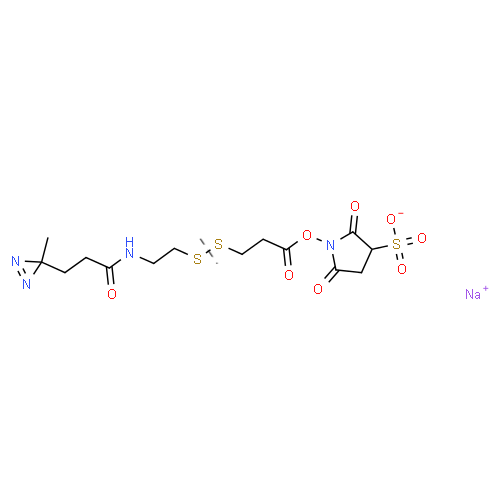 |
| AMAS | No | C_10_H_8_N_2_O_6_ | 252.03823598412 | C_10_H_8_N_2_O_6_ | 252.03823598412 | 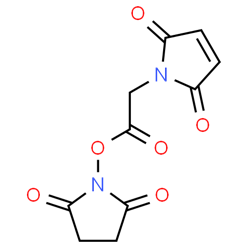 |
| BMPS | No | C_11_H_10_N_2_O_6_ | 266.05388604858 | C_11_H_10_N_2_O_6_ | 266.05388604858 | 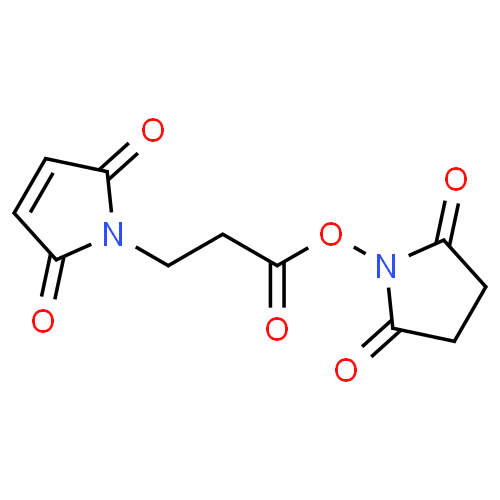 |
| EMCA | No | C_10_H_13_NO_4_ | 211.08445790169998 | C_10_H_13_NO_4_ | 211.08445790169998 | 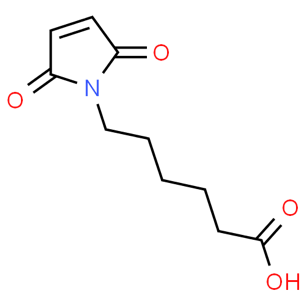 |
| EMCS | No | C_14_H_16_N_2_O_6_ | 308.10083624196 | C_14_H_16_N_2_O_6_ | 308.10083624196 | 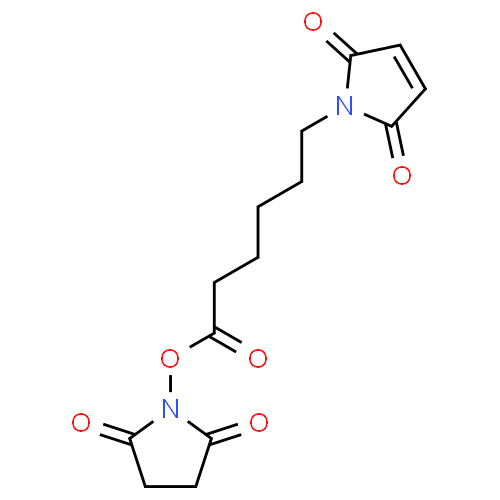 |
| GMBS | No | C_12_H_12_N_2_O_6_ | 280.06953611304004 | C_12_H_12_N_2_O_6_ | 280.06953611304004 | 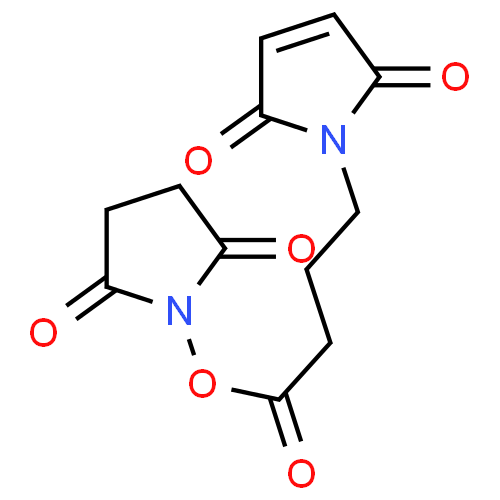 |
| LC-SMCC | No | C_22_H_29_N_3_O_7_ | 447.20055028495005 | C_22_H_29_N_3_O_7_ | 447.20055028495005 | 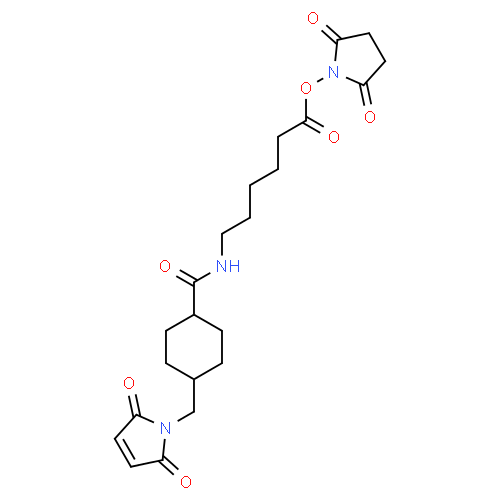 |
| LC-SPDP | Yes | C_18_H_23_N_3_O_5_S_2_ | 425.10791320122996 | C_13_H_19_N_2_O_5_S  C_5_H_4_NS | 315.10146789348  110.00644530775 | 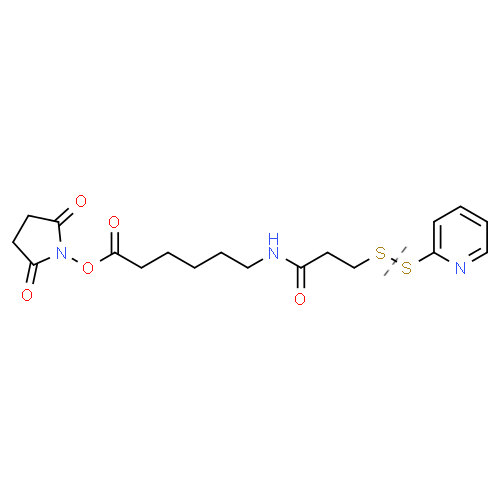 |
| MBS | No | C_15_H_10_N_2_O_6_ | 314.05388604858 | C_15_H_10_N_2_O_6_ | 314.05388604858 | 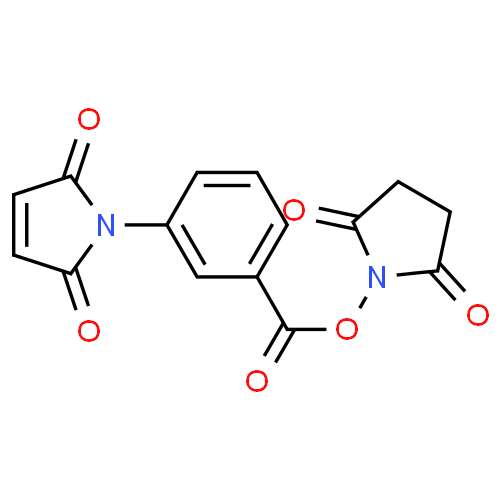 |
| PEG12-SPDP | Yes | C_39_H_65_N_3_O_17_S_2_ | 911.3755399897301 | C_5_H_4_NS  C_34_H_61_N_2_O_17_S | 110.00644530775  801.36909468198 | 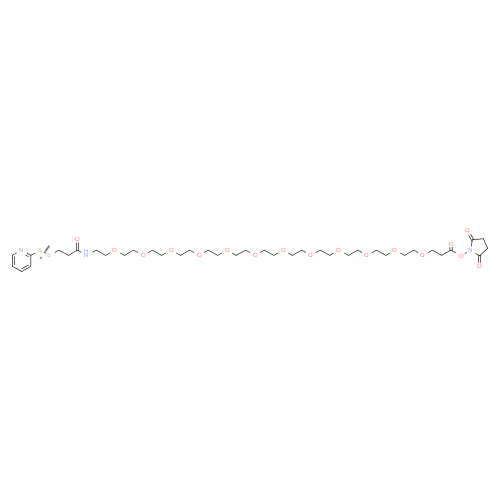 |
| PEG4-SPDP | Yes | C_23_H_33_N_3_O_9_S_2_ | 559.16582200181 | C_5_H_4_NS  C_18_H_29_N_2_O_9_S | 110.00644530775  449.15937669406 | 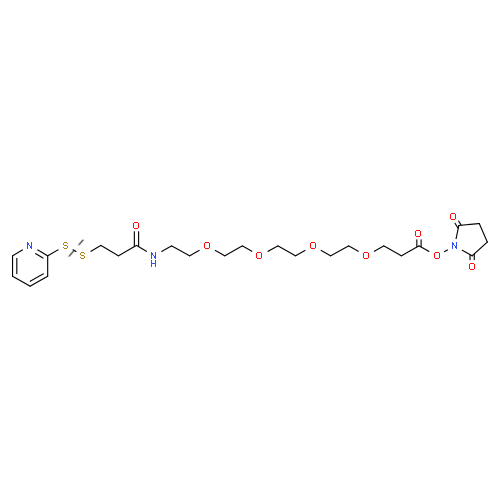 |
| SBAP | No | C_9_H_11_N_2_O_5_Br | 305.98513406124 | C_9_H_11_N_2_O_5_Br | 305.98513406124 | 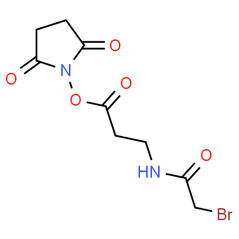 |
| SIA | No | C_6_H_6_INO_4_ | 282.93415457609 | C_6_H_6_INO_4_ | 282.93415457609 | 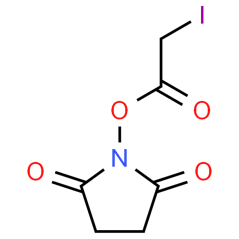 |
| SIAB | No | C_13_H_11_IN_2_O_5_ | 401.97126836124005 | C_13_H_11_IN_2_O_5_ | 401.97126836124005 | 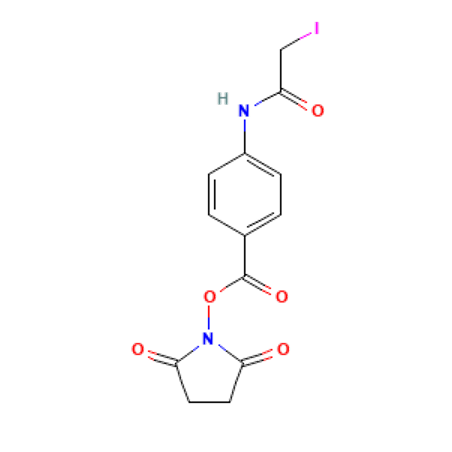 |
| SM(PEG)12 | No | C_38_H_63_N_3_O_19_ | 865.40557681561 | C_38_H_63_N_3_O_19_ | 865.40557681561 | 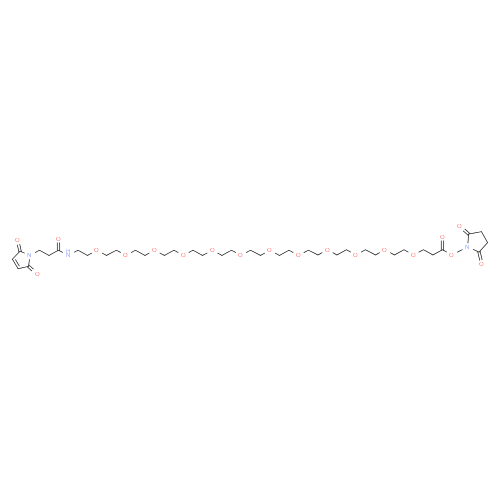 |
| SM(PEG)2 | No | C_18_H_23_N_3_O_9_ | 425.14342933071 | C_18_H_23_N_3_O_9_ | 425.14342933071 | 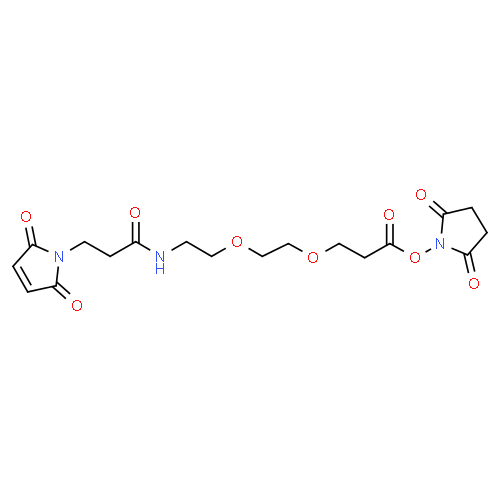 |
| SM(PEG)24 | No | C_62_H_111_N_3_O_31_ | 1393.72015379749 | C_62_H_111_N_3_O_31_ | 1393.72015379749 | 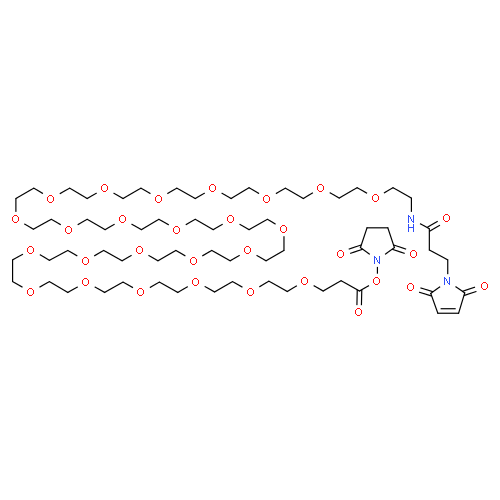 |
| SM(PEG)4 | No | C_22_H_31_N_3_O_11_ | 513.19585882769 | C_22_H_31_N_3_O_11_ | 513.19585882769 | 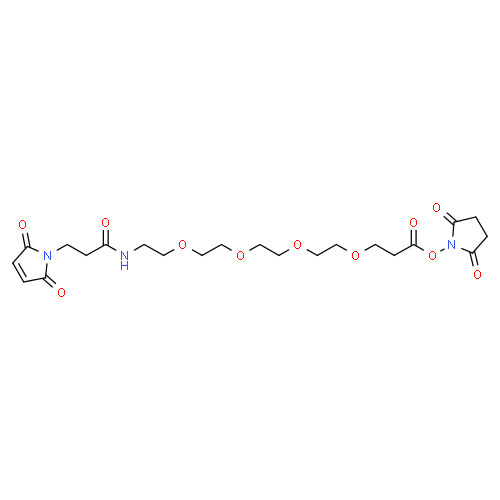 |
| SM(PEG)6 | No | C_26_H_39_N_3_O_13_ | 601.24828832467 | C_26_H_39_N_3_O_13_ | 601.24828832467 | 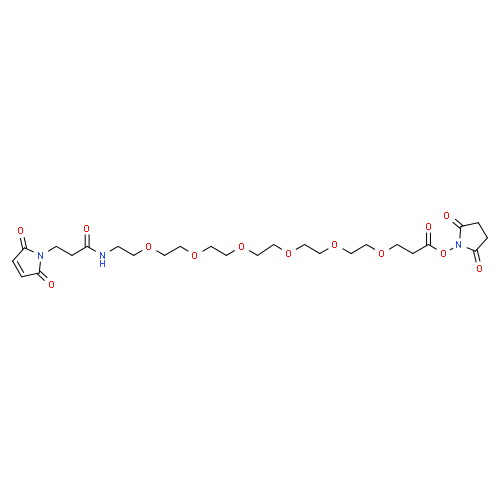 |
| SM(PEG)8 | No | C_30_H_47_N_3_O_15_ | 689.30071782165 | C_30_H_47_N_3_O_15_ | 689.30071782165 | 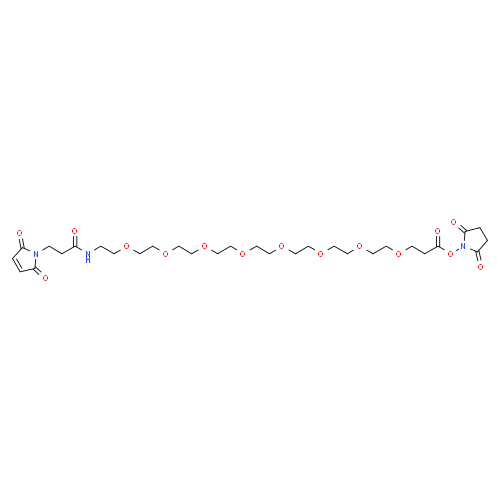 |
| SMCC | No | C_16_H_18_N_2_O_6_ | 334.11648630642003 | C_16_H_18_N_2_O_6_ | 334.11648630642003 | 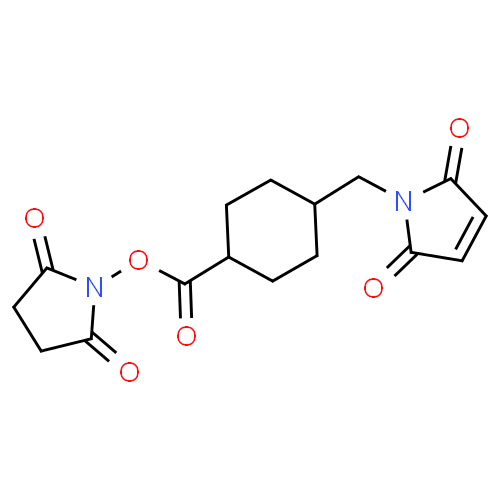 |
| SMPB | No | C_18_H_16_N_2_O_6_ | 356.10083624196 | C_18_H_16_N_2_O_6_ | 356.10083624196 | 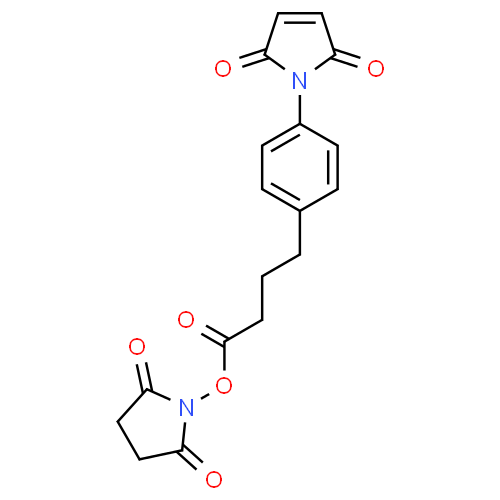 |
| SMPH | No | C_17_H_21_N_3_O_7_ | 379.13795002711004 | C_17_H_21_N_3_O_7_ | 379.13795002711004 | 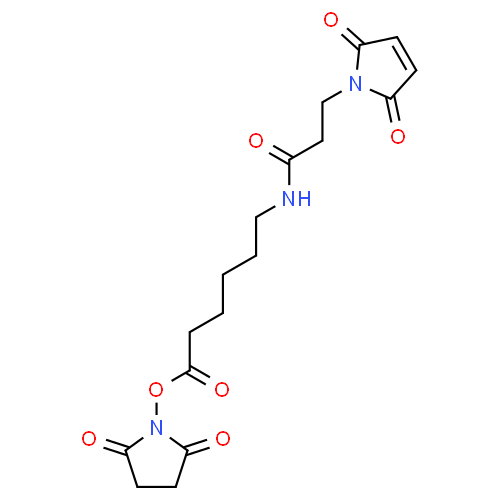 |
| SMPT | Yes | C_18_H_16_N_2_O_4_S_2_ | 388.05514935162 | C_5_H_4_NS  C_13_H_12_NO_4_S | 110.00644530775  278.04870404387003 | 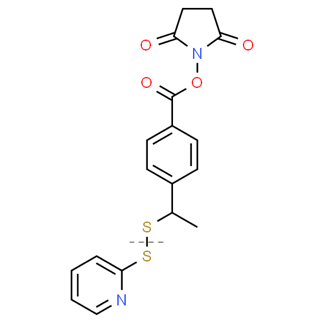 |
| SPDP | Yes | C_12_H_12_N_2_O_4_S_2_ | 312.02384922270005 | C_5_H_4_NS  C_7_H_8_NO_4_S | 110.00644530775  202.01740391495002 | 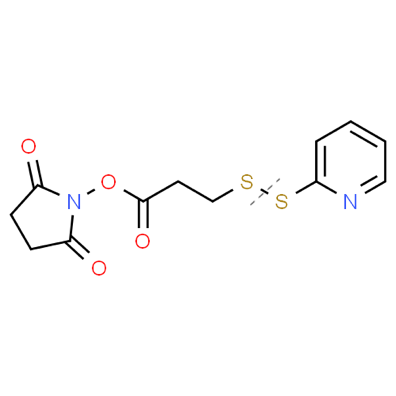 |
| Sulfo-EMCS | No | C_14_H_15_N_2_NaO_9_S | 410.03959552484 | C_14_H_15_N_2_NaO_9_S | 410.03959552484 | 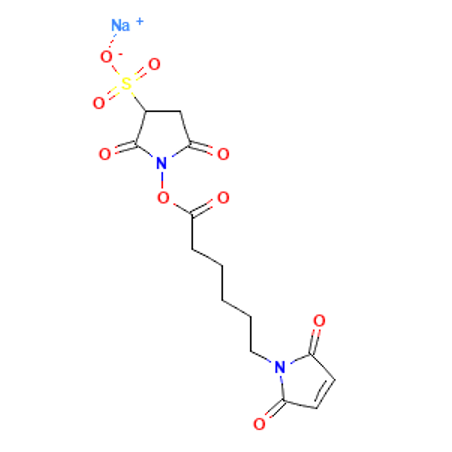 |
| Sulfo-GMBS | No | C_12_H_11_N_2_NaO_9_S | 382.00829539592 | C_12_H_11_N_2_NaO_9_S | 382.00829539592 | 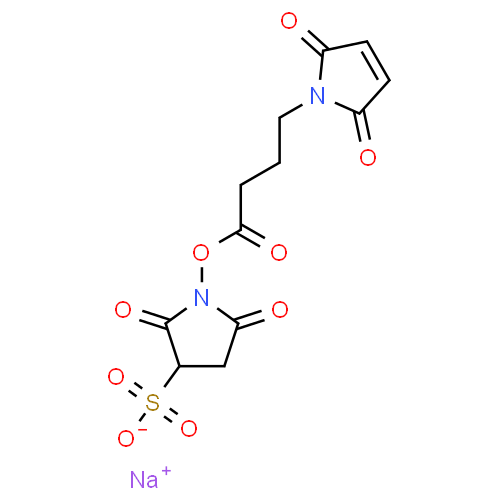 |
| Sulfo-KMUS | No | C_19_H_25_N_2_NaO_9_S | 480.11784584714 | C_19_H_25_N_2_NaO_9_S | 480.11784584714 | 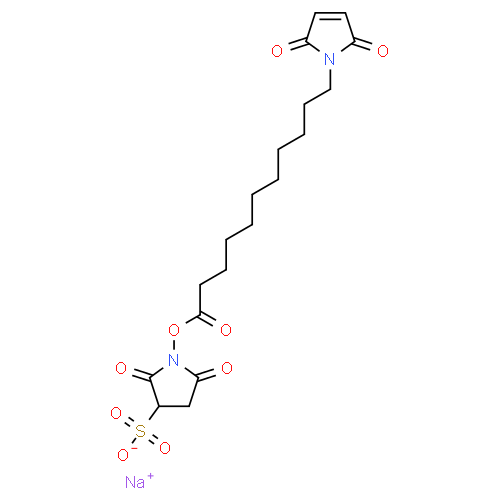 |
| Sulfo-LC-SPDP | Yes | C_18_H_22_N_3_NaO_8_S_3_ | 527.04667248411 | C_13_H_18_N_2_NaO_8_S_2_  C_5_H_4_NS | 417.0402271763601  110.00644530775 | 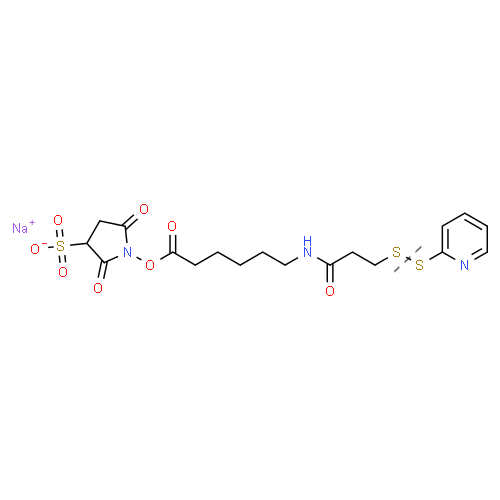 |
| Sulfo-MBS | No | C_15_H_9_N_2_NaO_9_S | 415.99264533146004 | C_15_H_9_N_2_NaO_9_S | 415.99264533146004 | 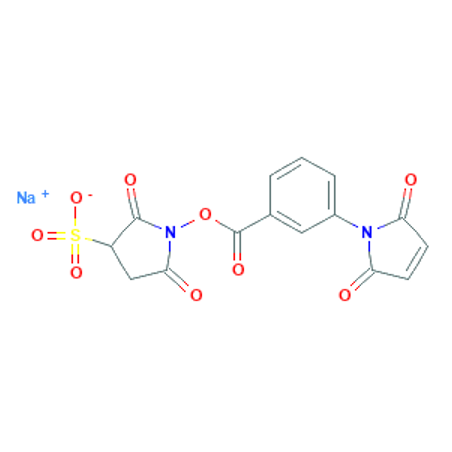 |
| Sulfo-SIAB | No | C_13_H_10_IN_2_NaO_8_S | 503.91002764412 | C_13_H_10_IN_2_NaO_8_S | 503.91002764412 | 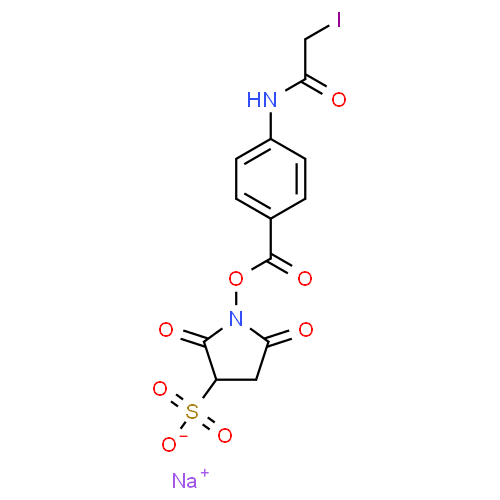 |
| Sulfo-SMCC | No | C_16_H_17_N_2_NaO_9_S | 436.0552455893 | C_16_H_17_N_2_NaO_9_S | 436.0552455893 | 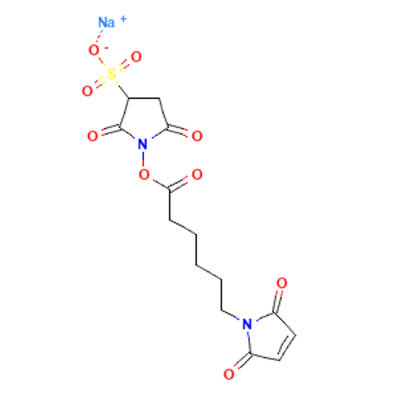 |
| Sulfo-SMPB | No | C_18_H_15_N_2_NaO_9_S | 458.03959552483997 | C_18_H_15_N_2_NaO_9_S | 458.03959552483997 | 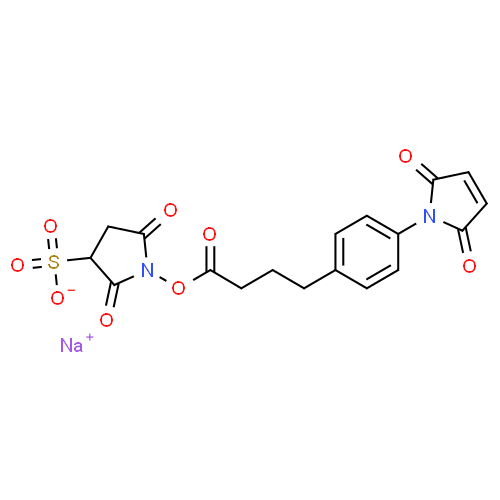 |
| IA-Alkyne | No | C_8_H_12_INO | 264.99636091076 | C_8_H_12_INO | 264.99636091076 | 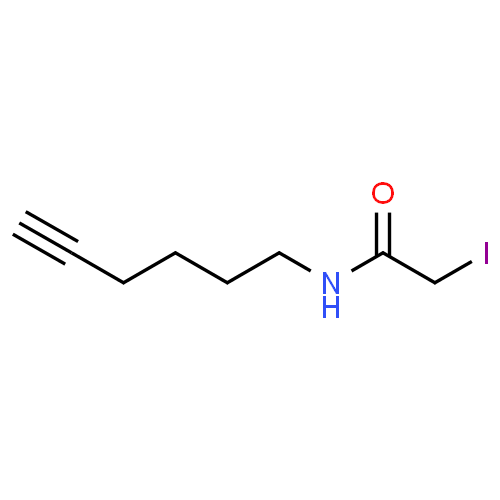 |
| DBCO-PEG4-maleimide | No | C_36_H_42_N_4_O_9_ | 674.2951789475101 | C_36_H_42_N_4_O_9_ | 674.2951789475101 | 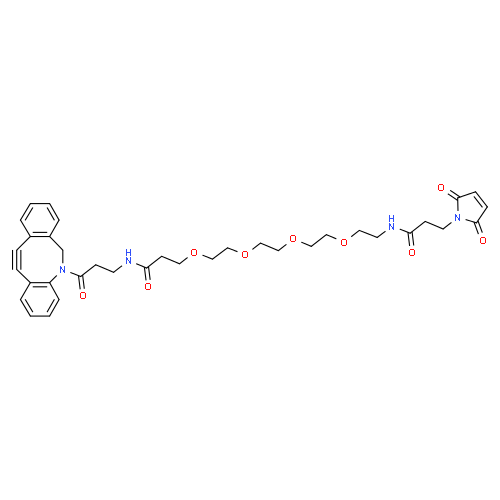 |
| MPBH | No | C_14_H_16_ClN_3_O_3_ | 309.08801906968006 | C_14_H_16_ClN_3_O_3_ | 309.08801906968006 | 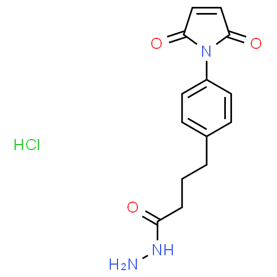 |
| PDPH | Yes | C_8_N_3_S_2_OH_11_ | 229.03435433619003 | C_5_NSH_4_  C_3_N_2_SOH_7_ | 110.00644530775  119.02790902843999 | 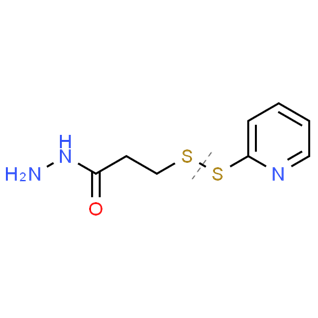 |
| PMPI | No | C_11_H_6_N_2_O_3_ | 214.03784206095003 | C_11_H_6_N_2_O_3_ | 214.03784206095003 | 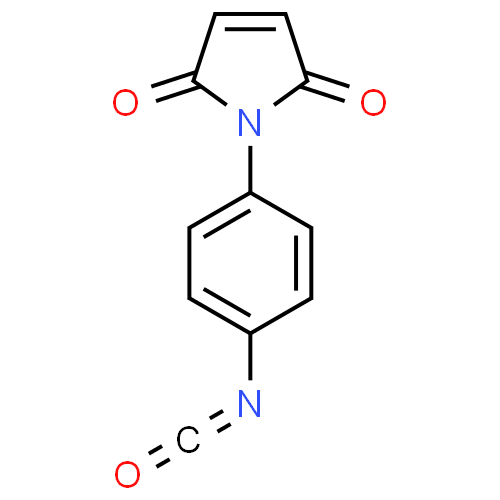 |
| BM(PEG)2 | No | C_14_H_16_N_2_O_6_ | 308.10083624196 | C_14_H_16_N_2_O_6_ | 308.10083624196 | 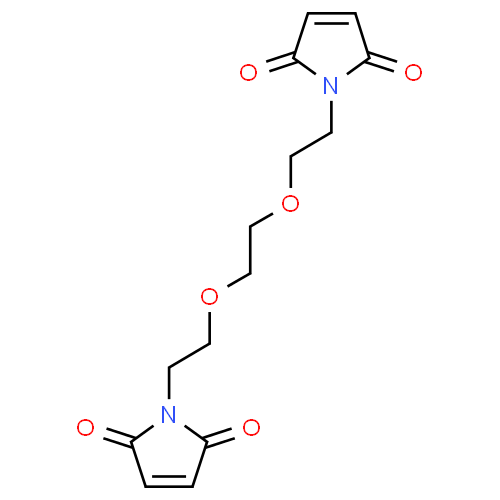 |
| BM(PEG)3 | No | C_16_H_20_N_2_O_7_ | 352.12705099045 | C_16_H_20_N_2_O_7_ | 352.12705099045 | 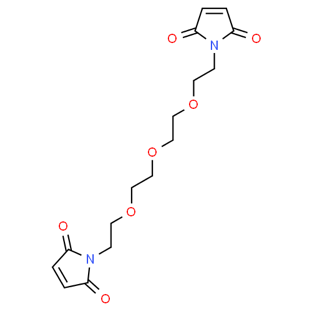 |
| BMB | No | C_12_H_12_N_2_O_4_ | 248.07970687390002 | C_12_H_12_N_2_O_4_ | 248.07970687390002 | 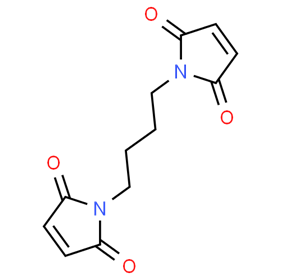 |
| BMH | No | C_14_H_16_N_2_O_4_ | 276.11100700282003 | C_14_H_16_N_2_O_4_ | 276.11100700282003 | 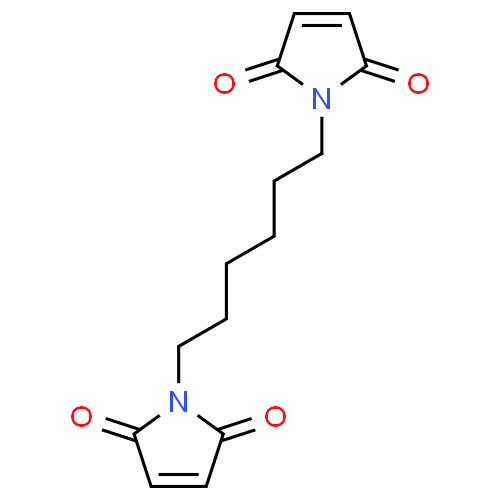 |
| BMOE | No | C_10_H_8_N_2_O_4_ | 220.04840674498 | C_10_H_8_N_2_O_4_ | 220.04840674498 | 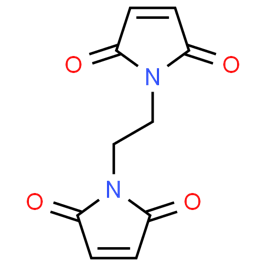 |
| DTME | Yes | C_12_H_12_N_2_S_2_O_4_ | 312.0238492227 | C_6_H_6_NSO_2_  C_6_H_6_NSO_2_ | 156.01192461135  156.01192461135 | 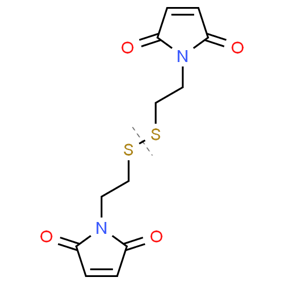 |
| TMEA | No | C_18_H_18_N_4_O_6_ | 386.12263431528004 | C_18_H_18_N_4_O_6_ | 386.12263431528004 | 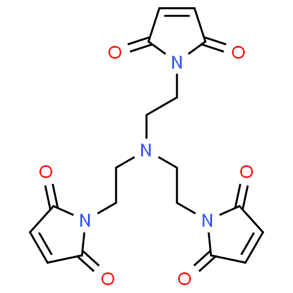 |
| SBA | No | C_6_H_6_BrNO_4_ | 234.94802027609 | C_6_H_6_BrNO_4_ | 234.94802027609 | 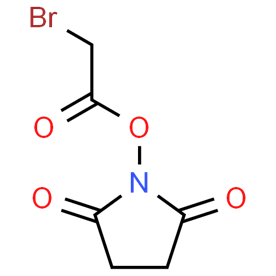 |
| DSSeb | No | C_18_H_24_N_2_O_8_ | 396.15326573894004 | C_18_H_24_N_2_O_8_ | 396.15326573894004 | 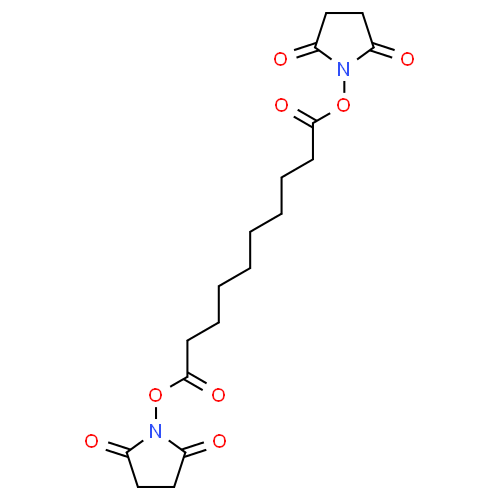 |
| Sulfo-SIA | No | C_6_H_7_INO_7_S | 363.89879464143 | C_6_H_7_INO_7_S | 363.89879464143 | 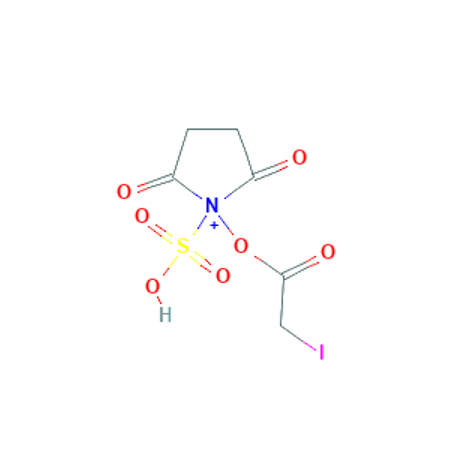 |
| SPDB | Yes | C_13_H_14_N_2_O_4_S_2_ | 326.03949928715997 | C_8_H_10_NO_4_S  C_5_H_4_NS | 216.03305397941  110.00644530775 | 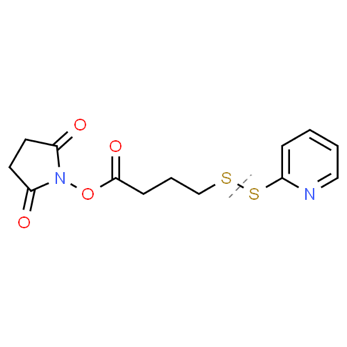 |
